# Inter- and intraspecific differences in *Drosophila* cold tolerance are linked to hindgut reabsorption capacity

**DOI:** 10.1101/774653

**Authors:** Mads Kuhlmann Andersen, Johannes Overgaard

## Abstract

Maintaining extracellular osmotic and ionic homeostasis is crucial to maintain organismal function. In insects, hemolymph volume and ion content is regulated by the combined actions of the secretory Malpighian tubules and reabsorptive hindgut. When exposed to stressful cold, homeostasis is gradually disrupted, characterized by a debilitating increase in extracellular K^+^ concentration (hyperkalemia). In accordance with this paradigm, studies have found a strong link between the cold tolerance of insect species and their ability to maintain ion and water homeostasis at low temperature. This is also the case for drosophilids where studies have already established how inter- and intra-specific differences in cold tolerance are linked to the secretory capacity of Malpighian tubules. However, presently there is little information on the effects of temperature on the reabsorptive capacity of the hindgut in *Drosophila.* To address this question we developed a novel method that allows for continued measurements of hindgut ion and fluid reabsorption in *Drosophila.* Firstly we demonstrate that this assay is temporally stable (> 3 hours) and that the preparation is responsive to humoral stimulation and pharmacological intervention of active and passive transport in accordance with the current insect hindgut reabsorption model. Using this method at benign (24°C) and low temperature (3°C) we investigated how cold acclimation or cold adaptation affected the thermal sensitivity of osmoregulatory function. We found that cold tolerant *Drosophila* species and cold-acclimated *D. melanogaster* are innately better at maintaining rates of fluid and Na^+^ reabsorption at low temperature. Furthermore, cold adaptation and acclimation causes a relative reduction in K^+^ reabsorption at low temperature. These characteristic responses of cold adapted/acclimated *Drosophila* will act to promote maintenance of ion and water homeostasis at low temperature and therefore provide further links between adaptations in osmoregulatory capacity of insects and their ability to tolerate cold exposure.

## Introduction

Maintenance of a relatively stable extracellular environment is essential for cell function and ultimately survival of most animals (Bernard, 1872). The concentrations of major ions and the volume of the extracellular fluid are maintained primarily through the actions of the secretory Malpighian tubules and the absorptive hindgut in insects (Beyenbach and Piermarini, 2008; Edney, 1977; Phillips, 1970). These osmoregulatory organs act in synchrony to balance the passive movement of ions and water across gut epithelia and integument whereby they help to secure ion and water homeostasis. Secretion and reabsorption of ions and water involve energy-demanding ion pumps that, accompanied by co-transporters and ion exchangers, regulate movement of ions and water (Beyenbach and Piermarini, 2011; O’Donnell et al., 2003; Phillips, 1981; Phillips, 1970). Active transporters are temperature-sensitive, and exposure to low temperature will therefore tend to reduce the capacity for secretion and reabsorption of ions (Gerber and Overgaard, 2018; MacMillan et al., 2015a; Yerushalmi et al., 2018). The loss of active transport can therefore ultimately lead to the characteristic, cold-induced loss of extracellular ion and water balance observed in chill-susceptible insects (Andersen et al., 2017b; Des Marteaux and Sinclair, 2016; Koštál et al., 2007; Koštál et al., 2004; Koštál et al., 2006; MacMillan et al., 2015a; MacMillan and Sinclair, 2011; Overgaard and MacMillan, 2017).

The loss of ion balance, particularly the increase in extracellular K^+^, is linked to chill injury (Andersen et al., 2017a; Bayley et al., 2018; MacMillan et al., 2015c; Overgaard and MacMillan, 2017) and can be ascribed to two main processes: 1) K^+^ leaks into the hemolymph *via* trans- or paracellular pathways faster than it can be removed (by the Malpighian tubules), or 2) Na^+^ and water leak into the gut faster than it can be reabsorbed resulting a loss of hemolymph volume which concentrates the K^+^ remaining in the hemolymph (MacMillan et al., 2015b; MacMillan and Sinclair, 2011; Overgaard and MacMillan, 2017). Consistent with this paradigm there are now several studies that have shown how cold adapted or cold acclimated insects are characterized by superior osmoregulatory function at low temperature (Andersen et al., 2017c; Des Marteaux et al., 2017; Gerber and Overgaard, 2018; MacMillan et al., 2015a; Yerushalmi et al., 2018).

The majority of studies investigating osmoregulatory organs in insects have used model species such as *Drosophila, Locusta* and *Rhodnius* and collectively these studies have been biased towards studies of epithelial transport in Malpighian tubules. This “bias” is likely influenced by the availability of the “Ramsay assay” which represents an easy and effective method to study epithelial transport and regulation in insects (see Ramsay (1954), but also Dow et al. (1994), Rheault and O’Donnell (2004), and Davies et al. (2019) for more recent applications). However, Malpighian tubules only represent the “secretory half’ of the osmoregulatory organs and historically there has been a shortage of studies investigating hindgut reabsorbtion in small model insects such as *Drosophila* (Black et al., 1987; Hanrahan et al., 1984; Phillips et al., 1987; Phillips et al., 1996). However, recent studies of osmoregulatory function of drosophila has become possible with the use of scanning ion-selective electrode technique (SIET) which estimates the net ion flux, but cannot quantify bulk ion or fluid transport in these small species (Andersen et al., 2017c; D’Silva et al., 2017; Donini and O’Donnell, 2005; Yerushalmi et al., 2018). Studies using SIET have already indicated that cold acclimation and adaptation influences ion and water reabsorption capacity in *Drosophila* (Andersen et al., 2017c; Yerushalmi et al., 2018) but considering the important role of the insect hindgut it is valuable to develop a system that can directly measure the combined actions of active and passive ion and water movements under *in vitro* conditions that mimic *in vivo* ion and temperature conditions.

The transport mechanisms currently known to occur in the hindgut (i.e. the rectal pads) of herbivorous insects include basolateral Na^+^/K^+^ ATPases which maintain a favorable gradient for Na^+^ reabsorption across the apical membrane aided by H^+^/Na^+^, amino acid/Na^+^, and NH_4_^+^/Na^+^ exchangers. Reabsorption of K^+^ is intimately linked to apical Cl^−^ transport, and although the exact mechanisms is still debated, two main non-mutually exclusive hypotheses suggest that Cl^¬^is reabsorbed *via* 1) H^+^ recycling through H^+^/Cl^−^ symporters and apical V-type H^+^ ATPase activity and/or 2) direct Cl^−^ reabsorption *via* an electrogenic Cl^−^ pump (see Gerencser and Zhang (2003), Hanrahan and Phillips (1983), Phillips (1981), Phillips et al. (1987), and Phillips et al. (1996)). Fluid reabsorption is thought to occur through the scalariform complex in the hindgut (Phillips, 1981; Wall and Oschman, 1975; Wall and Oschman, 1970). Here, the combination of a highly convoluted intercellular space and a high density of energy-demanding Na^+^/K^+^ ATPases in the intercellular membrane creates an osmotic gradient through the paracellular space of the rectal pads, drawing water through to the basolateral side of the epithelium (Gupta et al., 1980; Phillips et al., 1987). Despite this knowledge of transport mechanisms, few studies have actually measured the transport of fluid and ions across the hindgut of small insects and no study has so far been able to measure the *net* movement of ions and water across the *Drosophila* hindgut. It has therefore also been difficult to study the thermal dependency of net transport and evaluate putative role of hindgut capacity in relation to inter and intraspecific differences in cold tolerance of this species group.

In the present study, we present a novel assay capable of measuring *in vitro* ion and water reabsorption across the *Drosophila* hindgut. In our initial experiments we demonstrate the temporal stability of this assay and investigate how pharmacological blockade of central ion transporters and channels inhibit reabsorption. After these descriptive experiments we use the assay to compare hindgut reabsorption at benign (24°C) and low temperature (3°C) in three species of *Drosophila* (*D. montana*, *D*. *melanogaster* and *D*. *birchii*) characterized by large differences in cold tolerance. Similarly, we use this assay to evaluate the effects of cold acclimation by investigating hindgut transport in in cold- and warm-acclimated *Drosophila melanogaster.* With these experiments we test the hypothesis that cold tolerant species and cold-acclimated *D. melanogaster* exhibit adaptive changes in reabsorption capacity that will help them to prevent hemolymph hyperkalemia during cold exposure. Specifically, we hypothesize that cold acclimation and cold adaptation is associated with the ability to suppress K^+^ reabsorption and to maintain Na^+^ and water reabsorption during exposure to low temperature.

## Materials and Methods

### Animal husbandry

The present study investigated three species of *Drosophila* (*Drosophila montana* (Patterson, 1943), *Drosophila melanogaster* (Meigen, 1830), and *Drosophila birchii* (Dobzhansky and Mather, 1961), see **Table S1** for their origin) that are characterized by considerable differences in cold tolerance. Thus, *D. montana* is a cold tolerant species, *D. melanogaster* has intermediate tolerance and *D. birchii* is a cold sensitive species (Andersen et al., 2015; Andersen et al., 2017c; Kellermann et al., 2012; MacMillan et al., 2015a). All species were reared under common garden conditions at 19°C where they were fed on oatmeal-based Leeds medium (for one litre of water: 60 g yeast, 40 g sucrose, 30 g oat meal, 16 g agar, 12 g methyl paraben, and 1.2 mL acetic acid). Flies were kept in 200 mL bottles containing 40 mL of medium. To produce experimental flies, adult flies were allowed to oviposit for two hours to two days (depending on the species) in bottles with media to ensure a rearing density of 100-200 individuals per bottle. Newly emerged flies were transferred to vials (7 mL medium) and left to mature for 6-9 days before experiments.

For *D. melanogaster* flies we performed additional experiments where flies were acclimated at 15, 19 and 23°C, respectively. To produce these flies, adults were allowed to oviposit for 2 h at 19°C after which the egg-containing bottles were moved to thermal cabinets at 15, 19 or 23°C. Similar to the interspecific study the newly emerged flies were collected in vials and stored at their developmental temperature before females were used for experiments once they reached 6 to 9 days of age.

### Quantitative measurement of ion and water reabsorption in Drosophila hindguts

Individual flies were briefly submerged in 70% ethanol for sedation and females were then transferred to a glass Petri dish containing standard *Drosophila* saline with a layer of silicone elastomer at the bottom (Sylgaard 184, Dow Corning Corp., Midland, MI, USA) (saline: 137 mM Na^+^, 15 mM K^+^, 158.5 mM Cl^−^, 8.5 mM Mg^2+^, 2 mM Ca^2+^, 10.2 mM HCO_3_^−^, 4.3 mM H_2_PO_4_, 20 mM glucose, 10 mM glutamine, and 15 mM MOPS buffer, pH 7.0). The head, legs and wings were quickly removed, and the intact gut (crop, foregut, midgut, Malpighian tubules, and hindgut) was carefully dissected out, while ensuring that a small piece of cuticle remained around the anus (a representative drawing and a photo of the assay is shown in Fig. 1). The isolated gut was then gently transferred to a 50 μL droplet of *Drosophila* saline kept under paraffin oil. Using fine forceps (Dumont #5, 0.05 x 0.02 mm tips, Fine Science Tools Inc., Foster City, CA, USA) attached to a micromanipulator the hindgut was carefully drawn horizontally out of the droplet by gripping the small piece of cuticle still attached to the anus. This procedure isolates the hindgut from the remaining gut and Malpighian tubules (which remains immersed in the 50μL saline droplet). Any saline that still clings to the hindgut after this procedure is removed using a pulled glass capillary. Next, the isolated hindgut is superfused with a smaller droplet of saline (0.2 μL for *D. birchii* and *D. melanogaster* and 0.3 μL for *D. montana).* This droplet contains 100μM of amaranth so it is easy to see under the microscope and the amaranth also makes it easy to see if the small droplet (0.2-0.3 μL) fuses with the larger (50 μL) droplet. The small droplet clings to the hindgut because of surface tension, and will not envelop the anus because of the hydrophobic nature of the cuticle that still remains (Fig. 1). This preparation therefore has the added benefit that any discharge from the gut will be delivered outside of the saline droplet as fecal droplets which sink to the bottom of the petri dish.

**Figure 1.**
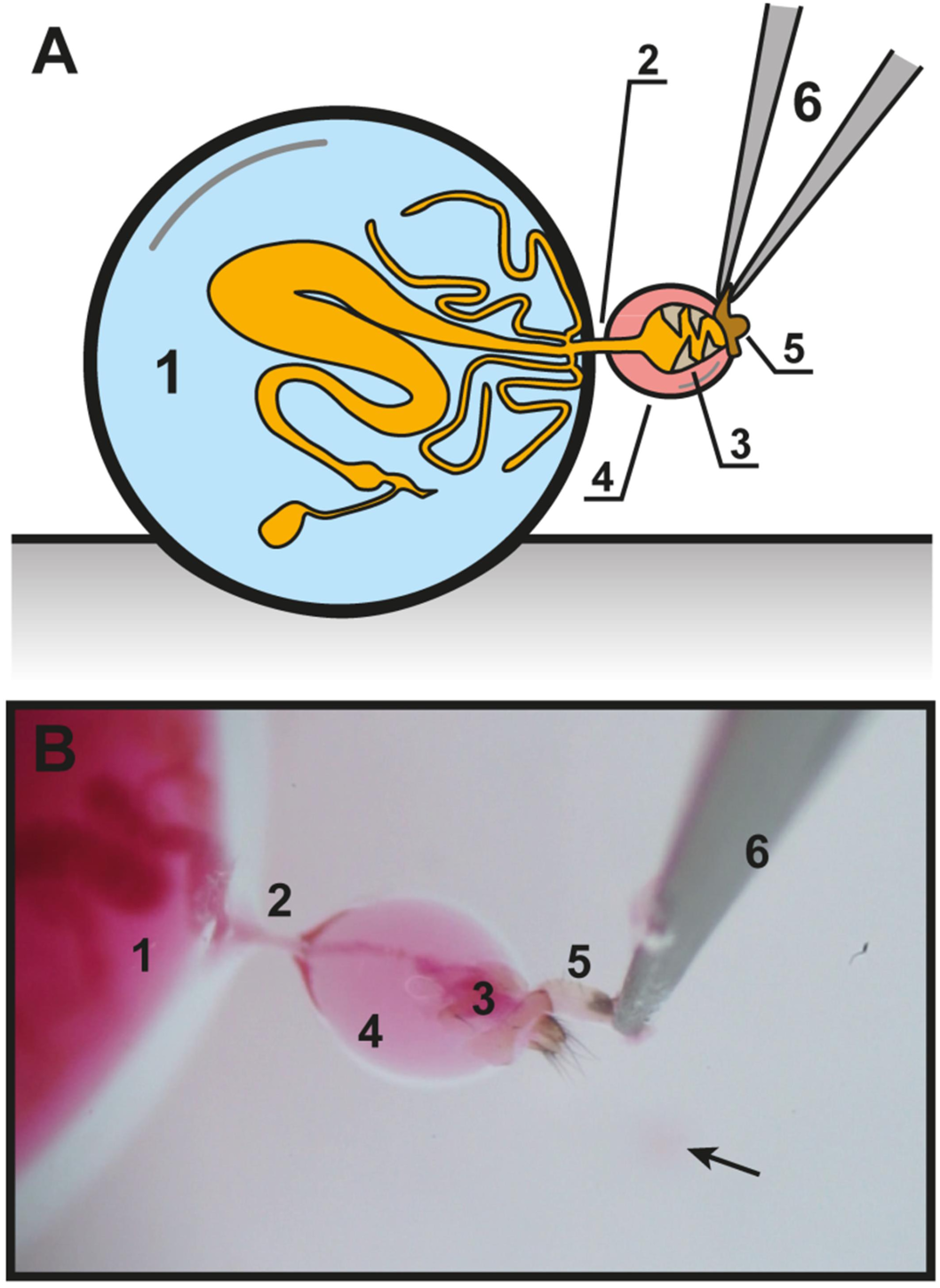
Schematic drawing and picture of the hindgut assay setup applied on Drosophila melanogaster. A) A schematic drawing of the hindgut assay and B) a picture of the assay applied to a specimen of *D. melanogaster* taken through a microscope. 1 = 50 μL droplet of standard *Drosophila* saline containing the fore- and midgut as well as the Malpighian tubules and most of the ileum, 2 = ileum, 3 = rectum with rectal pads, 4 = small droplet of standard *Drosophila* saline dyed with amaranth, 5 = anus with cuticle still attached, 6 = fine forceps. For the picture in panel B, where parts of the midgut and the Malpighian tubules are also visible, amaranth was added to the large droplet to help visualize the hindgut and to demonstrate that the primary urine and gut content moved through the ileum and into the hindgut to supply fluid and ions for reabsorption. Additionally, a small pink droplet of fecal matter can be seen in the background out of focus (arrow).

Over time the isolated hindgut will reabsorb ions and water and the small droplet surrounding the hindgut will gradually change its volume and ion-concentrations as the initial saline is altered by the reabsorbate. After a given time period (typically 1 h at 24°C) it is possible to sample the droplet and store it under the hydrated paraffin oil for later estimations of volume and ion concentration.

### Calculations of fluid and ion reabsorption

Estimates of fluid- and ion-reabsorption rates were derived from the measurements of changes in ion concentrations and volume of the droplet surrounding the hindgut. Volume of the mixture was estimated by imaging under a stereomicroscope (Carl Zeiss Stemi 2000-CS, Carl Zeiss A/S, Birkerød, DK) using a Sony α NEX 7 digital camera. After import into ImageJ (Schindelin et al., 2015), the diameter of the sample (*d*) was measured, and volume of the saline-reabsorbate mixture was calculated using the formula for volume of a sphere:

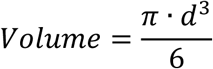

Water reabsorption rate was then calculated from the change in volume relative to the initial volume (0.2 or 0.3 μL) as follows:

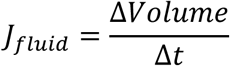

where *J*_*fluid*_ is the fluid reabsorption rate (nL min^−1^), *ΔVolume* is the change in volume (in nL), and *Δt* is the duration of the experiment (in minutes).

Ion concentrations (Na^+^ and K^+^) in the saline-reabsorbate mixture were measured using ion-sensitive microelectrodes that were constructed as described in MacMillan et al. (2015a). Before measurements the electrodes were calibrated in buffers with an order of magnitude difference in concentration (10 and 100 mM for K^+^, and 30 and 300 mM for Na^+^; osmolality was maintained by adding LiCl to the lower concentration standards). The recorded voltage change in these electrodes follows a Nernstian relationship with concentration (58.2 mV per 10-fold change in concentration). Only electrodes where a 10-fold change in concentration elicited a voltage-change between 52 and 62 mV (means ± standard deviation; K^+^: 56.3 ± 1.9 mV, Na^+^: 55.7 ± 1.7 mV) were deemed useful and outliers were discarded. Raw voltages were digitized using a MP100A data acquisition system and recorded using AcqKnowledge software (Biopac Systems, Goleta, CA, USA). Using these electrodes ion concentrations were calculated as follows:

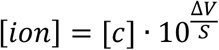

where [*ion*] is the concentration of either Na^+^ or K^+^ in the sample reabsorbate in mM, [c] is the concentration in the calibration buffer with the lowest calibration (30 or 10 mM, for Na^+^ and K^+^ respectively), *S* is the voltage-change observed with a 10-fold change in concentraiton, and ΔV is the difference between the voltage measured in the lower calibration fluid and the saline-reabsorbate mixture.

Measurement of ion reabsorption rate was calculated based on differences in ion concentrations and volume of the initial and final droplet surrounding the hindgut. To do this the following formula was used:

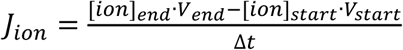

where *J*_*ion*_ is the rate of ion reabsorption (in pmol min^−1^), [*ion*]_end_ and [*ion*]_start_ are the concentrations of either Na^+^ or K^+^ in the saline-reabsorbate mixture at the experiment end and start, respectively (in mmol L^−1^). *V*_*start*_ is the volume of saline supplied to the hindgut in the beginning of the experiment (0.2 or 0.3 μL) and *V*_*end*_ is the volume of the saline-reabsorbate mixture at the end of the experiment. Δ*t* is the duration of the experiment (min).

### Experimental protocol and validation of assay

The primary scientific aim of the present study was to investigate if/how thermal adaptation and acclimation affects the hindgut capacity to maintain ion balance in *Drosophila* exposed to low temperature. To address this question a novel method was developed to assess quantitatively the movements of fluid and ions over the hindgut epithelia of *Drosophila.* To validate this new methodology a series of experiments was run to investigate the temporal stability of the assay and to briefly examine how active transport could be manipulated both at the hindgut and upstream at the level of the Malpighian tubule. These experiments were all performed in *D. melanogaster* at room temperature (22-23°C).

Initially the temporal stability of the assay was tested by measuring ion and fluid reabsorption rates every hour for four hours (N = 4). After ensuring the temporal stability of the assay, the effects of NaCN inhibition was investigated to examine if ion and water flux was related to active transport. These experiments were performed by initially measuring flux under control conditions over the first hour and subsequent measure flux following addition of 1 mmol L^−1^ NaCN (N = 5). NaCN was added to both the large and small droplet to block all aerobically driven active transport in the Malpighian Tubules, foregut and midgut (large droplet) as well as in the hindgut (small droplet). In two additional sets of experiments the effects of partial blockade of active transport was tested by adding NaCN (1 mmol L^−1^) to either the small droplet (to inhibit only the hindgut) or the large droplet (to inhibit only the upstream secretion) (N = 3 for each experiment).

To examine the role of major transporters involved in hindgut reabsorption the hindgut of *D. melanogaster* was subjected to three pharmacological agents that inhibit transporters and channels involved in ion and fluid reabsorption: 1) Na^+^/K^+^ ATPase activity was inhibited with ouabain (1 mmol L^−1^, N = 5); 2) V-type H^+^ ATPase activity was inhibited with bafilomycin A_1_ (10 μmol L^−1^ in 2% DMSO, N = 5) (a parallel experiment tested the effect of saline with 2% DMSO on hindgut reabsorption to ensure that this was not an effect of the DMSO (N = 3)); 3) general Cl^−^ transport was inhibited with 4,4’-diisothiocyano-2,2’-stilbenedisulfonic acid (DIDS, 100 μmol L^−1^, N = 4) (all chemicals from Sigma Aldrich, St. Louis, MO, USA, except for the bafilomycin A_1_ which was from Cayman Chemicals, Ann Arbor, MI, USA). For all treatments, the hindgut was first measured under control conditions after approx. 1 hour. The preparation was then incubated for 10-15 min with pharmacological agents before the small droplet was changed to assess the effects of pharmacological blockade over a subsequent hour (during this time the preparation was also exposed to the pharmacological agents).

We also investigated stimulation of active transport by stimulating the cAMP pathway. To do so, 1 mmol L^−1^ of the cAMP stimulator 8-bromoadenosine 3’, 5’-cyclic monophosphate (Sigma Aldrich, St. Louis, MO, USA) was added to the buffer (both droplets) to stimulate both secretion by the Malpighian tubules and reabsorption at the hindgut. Lastly, an experiment was performed in which the concentration of Na^+^ in the saline was lowered from the initial 137 mM to 65 mM (N = 6). For these experiments it was not possible to change the buffer in which the gut was suspended and these experiments are therefore not compared to the control situation of the same preparations. Instead the absolute transport rates are compared to the average of the other controls performed at room temperature (22-23°C).

### The effects of adaptation and thermal acclimation on hindgut reabsorption capacity

As mentioned above a main objective of this study was to investigate and compare the osmoregulatory capacity at benign and low temperature in three *Drosophila* species with marked differences in cold tolerance (N = 7 per species and temperature). Likewise, *D. melanogaster* acclimated to 15, 19 and 23°C (N = 7-9 per acclimation temperature and test temperature) were compared to investigate how acclimation influenced hindgut ion and water flux. For all these experiments ion and water flux was measured at room temperature (24°C) first, after which temperature of the preparation was lowered to 3°C by circulating a cooled 1: 1 mixture of water and ethylene-glycol through the water-jacketed stage, in which the preparation was kept. Once at 3°C, the procedure of preparing the assay was repeated to obtain repeated measurements of fluid and ion reabsorption. As expected the transport rates were much lower at low temperature and consequently measurements took longer to achieve a reasonable change in droplet volume and content (~ 3-5 h at 3°C compared to ~ 1 h at 24°C).

### Data analysis

To estimate the effect of temperature on the transport of ions and water the temperature coefficients (Q_10_) were calculated for all three species and for warm and cold acclimated *D. melanogaster* as follows:

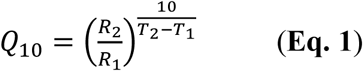

where R_2_ is the rate of reabsorption at the highest temperature (T_2_, 24°C) and R_1_ is the rate of reabsorption at the lowest temperature (T_1_, 3°C).

To examine if temperature influenced the selectivity of ion transport (Na^+^ transport relative to K^+^ transport), the temperature effect ratio was calculated as follows:

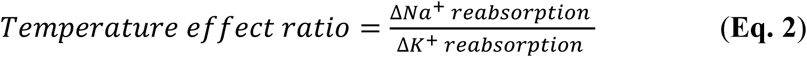

where the ΔNa^+^ reabsorption and ΔK^+^ reabsorption represent the absolute changes in Na^+^ and K^+^ reabsorption induced by exposure to 3°C relative to 24°C, respectively.

### Statistical analyses

All statistical analyses were performed in R 3.6.1 software (R Core Team, 2019). The stability of fluid and ion reabsorption over time was analyzed using a linear mixed effect model by using the lme() function in the nlme package for R (with time as a fixed factor and the fly as a random factor) followed by Dunnett’s *post hoc* tests to determine when rates of reabsorption deviated from control values. The effects of pharmacological interventions (NaCN inhibition, cAMP stimulation, inhibition with ouabain, bafilomycin A_1_, and DIDS, along with the effect of DMSO) on rates of reabsorption were analyzed using paired t-tests. The effect of lowering the Na^+^ concentration in the saline was tested using an unpaired Students t-test to comparing the “low Na^+^” reabsorption rate with the average of reabsorption rate from all other control measurements at room temperature that had been performed in “high Na^+^” saline.

The effects of species/acclimation and temperature on fluid and ion (Na^+^ and K^+^) reabsorption rates were analyzed using linear mixed effect models where species (*D. birchii, D. melanogaster* or *D*. *montana*) or acclimation temperature (15, 19 or 23°C) were included in the model as fixed factors along with temperature, while the individual fly was included as a random factor. Interspecific differences in Q_10_’s and temperature effect ratios were analyzed using separate one-way ANOVA’s and followed by Tukey’s Honest Significant Differences *post hoc* tests. It was, however, not possible to calculate Q_10_ for Na^+^ reabsorption of *D. birchii* (because of negative rates of reabsorption at low temperature) and the difference between *D. montana* and *D. melanogaster* was therefore analyzed using a Students t-test. A similar set of analyses were performed for the acclimation experiments. In all analyses, the critical level of significance was 0.05, and all values presented are means ± s.e.m. Full details of all analyses can be found in **Tables S2 and S3**.

## Results

### Experimental evaluation of hindgut assay

To examine the temporal stability of reabsorption rates in our novel hindgut assay, we collected samples every hour for 4 hours from hindguts of *D. melanogaster* operating at room temperature (Table 1). Fluid reabsorption showed a slow decline over time (P = 0.020), but remained stable for the two first hours before the decline became statistically significant. A similar temporal decline was found for Na^+^ reabsorption (P = 0.005), while the trend for a decline in K^+^ reabsorption did not reach statistical significance (P = 0.122). In the remaining experiments evaluating the effects of various pharmacological interventions we obtained a control measurement during the first hour followed by a measurement under “treatment conditions” during the following hour. The relative changes in activity associated with the pharmacological intervention can therefore be compared to the small (and non-significant) changes observed between hour 0 and 1 in the control for time (Table 1).

**Table 1.**
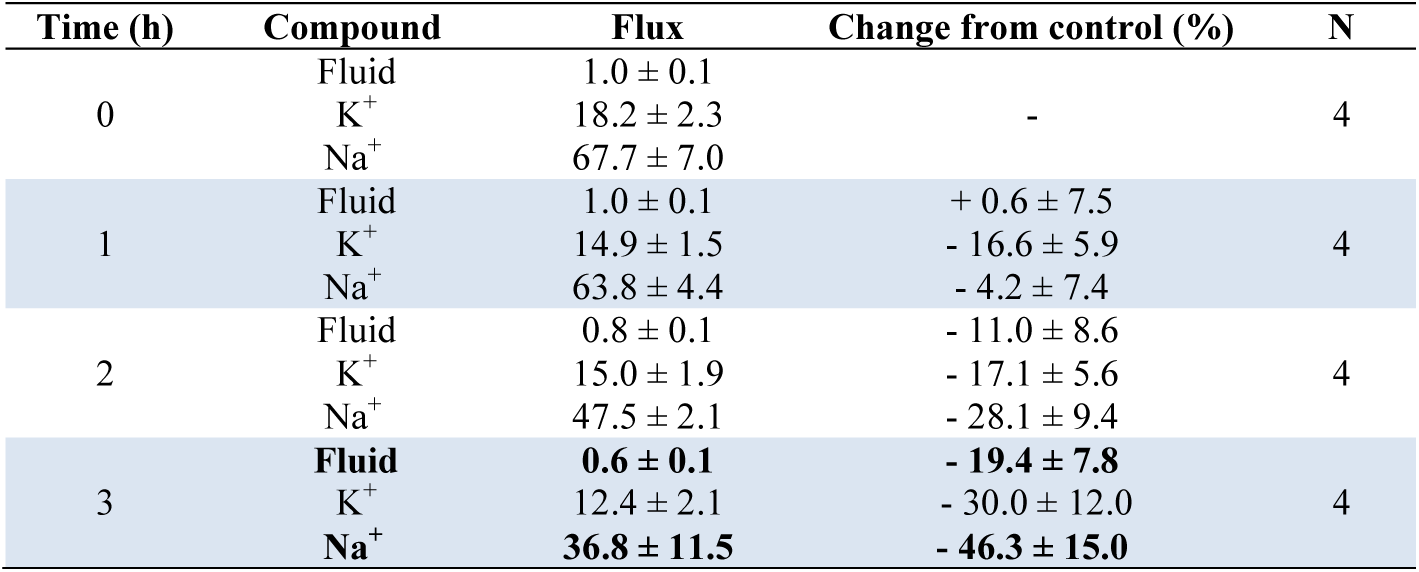
Stability over time of reabsorption of fluid and ions in the hindgut preparation measured in *D. melanogaster*. Fluid and ion reabsorption (in nL min^−1^ and pmol min^−1^, respectively) were measured every hour for three hours after a control measurement. Values presented in bold are significantly different from control values (based on separate Dunnett’s *post hoc* tests).

| Time (h) | Compound | Flux | Change from control (%) | N |
| --- | --- | --- | --- | --- |
| 0 | Fluid | 1.0 ± 0.1 | - | 4 |
|  | K<sup>+</sup> | 18.2 ± 2.3 |  |  |
|  | Na<sup>+</sup> | 67.7 ± 7.0 |  |  |
| 1 | Fluid | 1.0 ± 0.1 | + 0.6 ± 7.5 | 4 |
|  | K<sup>+</sup> | 14.9 ± 1.5 | - 16.6 ± 5.9 |  |
|  | Na<sup>+</sup> | 63.8 ± 4.4 | - 4.2 ± 7.4 |  |
| 2 | Fluid | 0.8 ± 0.1 | - 11.0 ± 8.6 | 4 |
|  | K<sup>+</sup> | 15.0 ± 1.9 | - 17.1 ± 5.6 |  |
|  | Na<sup>+</sup> | 47.5 ± 2.1 | - 28.1 ± 9.4 |  |
| 3 | **Fluid** | **0.6 ± 0.1** | **- 19.4 ± 7.8** | 4 |
|  | K<sup>+</sup> | 12.4 ± 2.1 | - 30.0 ± 12.0 |  |
|  | Na<sup>+</sup> | **36.8 ± 11.5** | **- 46.3 ± 15.0** |  |

Table 2 reports the control values from the preparations used in different experimental series and we found that the preparations were performing at stable rates across the different experiments. Thus, average fluid reabsorption rates were 1.0 nL min^−1^ (standard deviation: 0.1, range: 0.8 – 1.2), average K^+^ reabsorption rates were 17.2 pmol min^−1^ (standard deviation: 1.5, range 14.9 – 20.3), and average Na^+^ reabsorption rates were 69.3 pmol min^−1^ (standard deviation: 6.4, range 58.5 – 77.8).

**Table 2.** Results of pharmacological interventions on the reabsorption of fluid and ions in the hindgut of *D. melanogaster*. Here, control measurements were followed by an experiment in which the hindgut, the upstream fore- and midgut and Malpighian tubules, or both systems were exposed to a pharmacological agent to test the involvement of active transport and specific transporters in determining hindgut fluid and ion reabsorption. Fluid reabsorption is presented as nL min^−1^ while ion reabsorption (K^+^ and Na^+^) is in pmol min^−1^. Reabsorption rates presented in bold showed a significant effect of the intervention; for full statistical analyses **see Table S1**.

| Treatment | Applied to | Compound | Control | Treatment | N | P-value |
| --- | --- | --- | --- | --- | --- | --- |
| 1 mmol L<sup>-1</sup> NaCN | Hindgut | **Fluid** | **1.0 ± 0.0** | **0.4 ± 0.1** | 3 | **0.018** |
|  |  | K<sup>+</sup> | 16.2 ± 3.5 | 5.2 ± 1.8 |  | 0.055 |
|  |  | **Na<sup>+</sup>** | **58.5 ± 10.2** | **17.8 ± 8.7** |  | **0.014** |
| 1 mmol L<sup>-1</sup> NaCN | Malpighian tubules, fore- and midgut | **Fluid** | **1.2 ± 0.2** | **0.4 ± 0.1** | 3 | **0.005** |
|  |  | K<sup>+</sup> | **20.3 ± 5.3** | **8.1 ± 2.6** |  | **0.048** |
|  |  | **Na<sup>+</sup>** | **77.8 ± 17.5** | **23.1 ± 12.2** |  | **0.014** |
| 1 mmol L<sup>-1</sup> NaCN | Hindgut, Malpighian tubules, fore- and midgut | **Fluid** | **1.0 ± 0.1** | **0.2 ± 0.1** | 5 | **0.001** |
|  |  | K<sup>+</sup> | **17.3 ± 3.2** | **3.9 ± 1.0** |  | **0.004** |
|  |  | **Na<sup>+</sup>** | **71.1 ± 11.2** | **3.3 ± 4.3** |  | **0.002** |
| 1 mmol L<sup>-1</sup> ouabain | Hindgut | **Fluid** | **0.9 ± 0.1** | **0.4 ± 0.1** | 5 | **0.029** |
|  |  | K<sup>+</sup> | 16.4 ± 3.4 | 14.0 ± 2.9 |  | 0.265 |
|  |  | Na<sup>+</sup> | 73.0 ± 7.3 | 43.4 ± 8.0 |  | 0.058 |
| 10 µmol L<sup>-1</sup> bafilomycin A<sub>1</sub> (2% DMSO) | Hindgut | **Fluid** | **1.1 ± 0.1** | **0.6 ± 0.0** | 5 | **0.004** |
|  |  | K<sup>+</sup> | 18.2 ± 2.4 | 14.3 ± 2.5 |  | 0.350 |
|  |  | **Na<sup>+</sup>** | **70.8 ± 8.3** | **26.9 ± 3.4** |  | **0.006** |
| 2% DMSO | Hindgut | Fluid | 1.0 ± 0.1 | 0.9 ± 0.1 | 3 | 0.070 |
|  |  | K<sup>+</sup> | 14.9 ± 1.4 | 16.4 ± 1.4 |  | 0.081 |
|  |  | Na<sup>+</sup> | 72.7 ± 6.5 | 64.6 ± 3.0 |  | 0.237 |
| 100 µmol L<sup>-1</sup> DIDS | Hindgut | **Fluid** | **0.9 ± 0.1** | **0.1 ± 0.0** | 4 | **0.003** |
|  |  | K<sup>+</sup> | **17.6 ± 0.6** | **3.9 ± 1.1** |  | **0.002** |
|  |  | **Na<sup>+</sup>** | **71.9 ± 4.7** | **5.6 ± 2.1** |  | **0.001** |
| 1 mmol L<sup>-1</sup> 8-bromo-cAMP | Hindgut, Malpighian tubules, fore- and midgut | **Fluid** | **0.8 ± 0.1** | **1.1 ± 0.2** | 8 | **0.002** |
|  |  | K<sup>+</sup> | **16.8 ± 1.6** | **32.2 ± 3.7** |  | **0.002** |
|  |  | **Na<sup>+</sup>** | **59.5 ± 5.1** | **97.1 ± 8.8** |  | **0.002** |
| 65 mmol L<sup>-1</sup> Na<sup>+</sup> in saline | Hindgut, Malpighian tubules, fore- and midgut | Fluid | 1.0 ± 0.0* | 0.8 ± 0.1 | 6 | 0.091 |
|  |  | K<sup>+</sup> | 17.2 ± 0.1* | 17.2 ± 0.1 |  | 0.984 |
|  |  | **Na<sup>+</sup>** | **68.0 ± 3.0\*** | **37.6 ± 5.0** |  | **< 0.001** |

\*The control values for reabsorption in the experiments involving lowered [Na<sup>+</sup>] in the saline consist of 36 measurements performed in the other experiments under the same conditions (i.e. control measurements for all other experiments presented in this table and the control measurements from the time control).

The effects of pharmacological stimulation/inhibition of ion transport are also reported in Table 2 (see **Table S2** for statistical analyses). Hindgut reabsorption is clearly dependent on aerobic ATP production since exposure to 1 mmol L^−1^ NaCN caused a 59.5 ± 8.6 *%* reduction in fluid reabsorption rate and a 67.2 ± 7.5 % and 71.7 ± 9.5 % reduction in K^+^ and Na^+^ reabsorption rates, respectively. Hindgut reabsorption of fluid and ions is naturally also dependent on the provision of fluid and ions originating from the fore- and midgut as well as the Malpighian tubules. Thus, hindgut reabsorption was also reduced markedly (and significantly) when active transport was blocked upstream of the hindgut by using 1 mmol L^−1^ NaCN in the large droplet: Fluid reabsorption was reduced by 68.7 ± 7.0 %, K^+^ reabsorption by 61.9 ± 4.4 %, while Na^+^ reabsorption was reduced by 74.7 ± 10.0 %. In accordance with these results we found that 1 mmol L^−1^ NaCN exposure to both the hindgut and the upstream osmoregulatory organs resulted in a severe inhibition of fluid (−83.4 ± 6.4 %), K^+^ (−78.3 ± 2.3 %), and Na^+^ (−96.9 ± 4.9 %) reabsorption (Table 2).

We also briefly examined the role of major ion transporters and channels suspected to be involved in hindgut reabsorption of *Drosophila.* We first blocked Na^+^/K^+^ ATPase activity with ouabain (1 mmol L^−1^). This resulted in a 51.8 ± 13.6 % decrease in fluid reabsorption. K^+^ reabsorption remained relatively unchanged (2.0 ± 18.9 % decrease) while Na^+^ reabsorption showed a non-significant drop of 38.0 ± 11.7 %.

Application of the V-type H^+^ ATPase blocker bafilomycin A_1_ (10 ¼.mol L^−1^) resulted in a reduction of fluid reabsorption by 45.2 ± 4.7 %. K^+^ reabsorption was slightly, and non¬significantly reduced (17.0 ± 16.6 % decrease) while Na^+^ reabsorption decreased by 60.2 ± 5.7 %. The addition of bafilomycin A_1_ required the use of a 2% DMSO concentration in the saline. However, a parallel set of experiments showed that 2% DMSO alone had neither major, nor significant effects on reabsorption rates (Table 2).

DIDS (100 μ.mol L^−1^) was used to block all Cl^−^-related transporters and channels in the hindgut and the application of DIDS resulted in an almost complete abolishment of all transport; fluid reabsorption was reduced by 83.5 ± 3.9 %, K^+^ reabsorption by 77.7 ± 6.7 %, and Na^+^ reabsorption by 92.3 ± 2.8.

To examine if hindgut reabsorption is regulated through a cAMP dependent pathway the cAMP stimulator 8-bromo cAMP was added (1 mmol L^−1^) resulting in a 47.1 ± 10.3 % increase in fluid reabsorption associated, a doubling of K^+^ reabsorption (101.8 ± 28.1 % increase) and a 67.6 ± 18.7 % increase in Na^+^ reabsorption.

Finally, we examined the effects of lowering the Na^+^ concentration in the fluid surrounding both the upstream osmoregulatory organs (Malpighian tubule and fore-, and midgut) and the hindgut. Lowering the saline Na^+^ concentration from 137 mmol L^−1^ to 65 mmol L^−1^ elicited a small and non-significant reduction in fluid reabsorption (decreased by 19.1 ± 10.3 %). K^+^ reabsorption remained unchanged (−0.3 ± 6.9 %) while Na^+^ reabsorption was significantly reduced by 44.8 ± 7.3 %.

### Interspecific differences in hindgut transport during cold exposure

Hindgut fluid and ion reabsorption was measured at warm (24°C) and low (3°C) temperature in three *Drosophila* species characterized by high (*D. montana*), intermediate (*D. melanogaster*) and low (*D. birchii*) tolerance to cold. Fluid reabsorption rates were similar at warm temperature in all species (approximately ~ 0.9 nL min^−1^) (P = 0.716) (Fig. 2A). As hypothesized, fluid reabsorption decreased in all species during cold exposure (P < 0.001), and this effect tended to be more extreme in the most cold sensitive species such that fluid reabsorption rate was decreased to 0.06 ± 0.02 nL min^−1^ in *D. birchii*, 0.15 ± 0.03 nL min^−1^ in *D. melanogaster*, and 0.22 ± 0.04 in *D. montana*, despite not being statistically significant (P = 0.279). This tendency was more obvious (and significant, P = 0.004) when the response to lowered temperature was analyzed from the species specific Q_10_’s which ranged from 4.5 ± 0.8 for *D. birchii* to 2.6 ± 0.3 for *D. melanogaster* and 1.9 ± 0.2 for *D. montana* (Fig. 2B).

**Figure 2.**
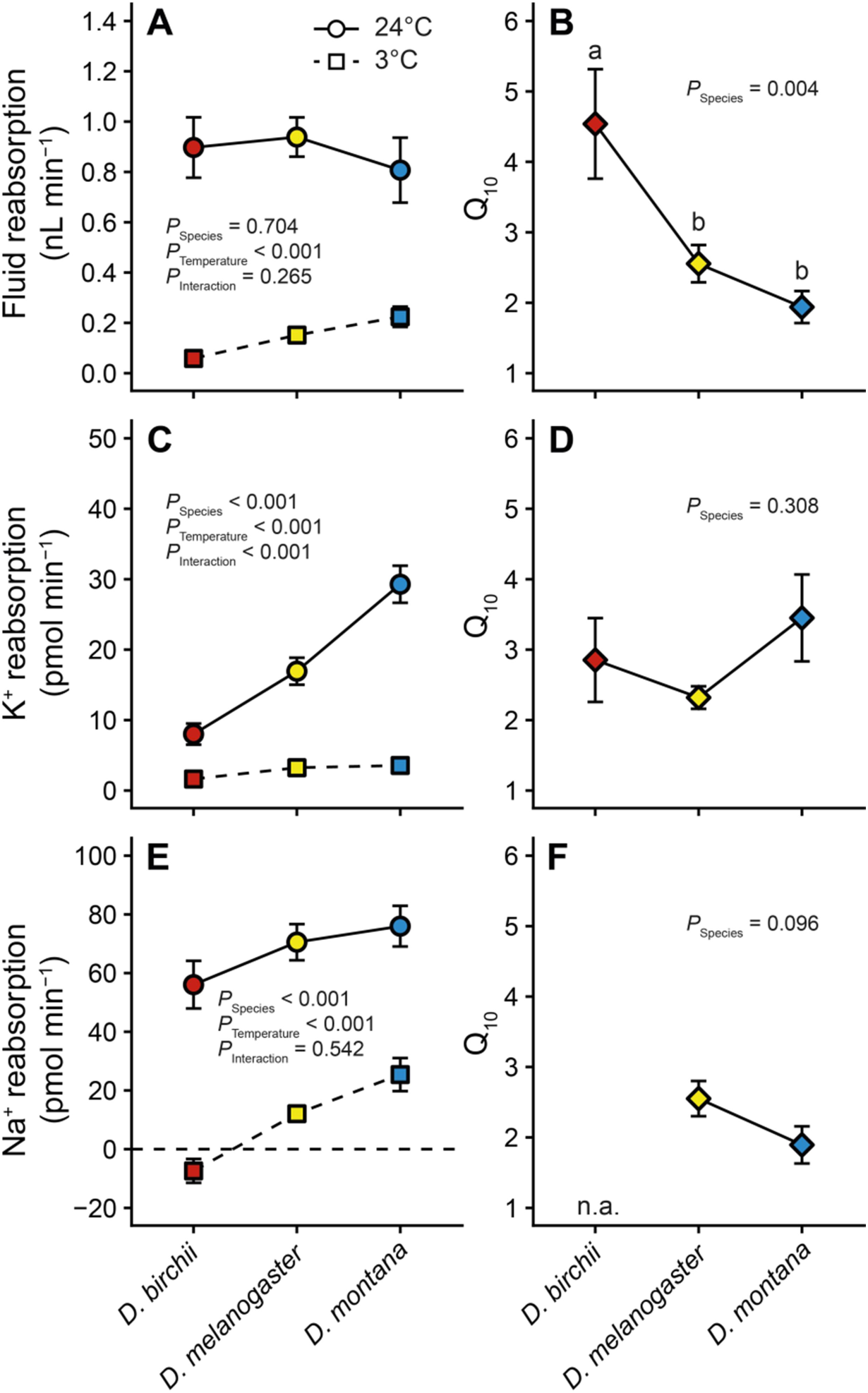
The effect of temperature on the fluid and ion reabsorption at the hindgut of three different Drosophila species. Hindgut reabsorption (left column) was measured at 24°C (circles) and 3°C (squares) and from these results the representative Q_10_’s were calculated (right column) for fluid (A and B), K^+^ (C and D), and Na^+^ (E and F). The horizontal, dashed line in panel C indicates zero reabsorption, and negative values indicate the transition to net secretion. N = 7 per species and temperature combination, and error bars not shown are covered by the symbols. For Na^+^ reabsorption, the switch to Na^+^ secretion prevented calculation of Q_10_.

The K^+^ reabsorption rate differed between the three species at 24°C with the highest rate (29.3 ± 2.6 pmol min^−1^) in the cold tolerant *D. montana*, an intermediate rate (16.9 ± 1.9 pmol min^−1^) in *D. melanogaster* and the lowest rate (8.0 ± 1.5 pmol min^−1^) in the cold sensitive *D. birchii* (P < 0.001) (Fig. 2C). Cooling to 3°C reduced K^+^ reabsorption in all species (P < 0.001), but did so in a species-specific manner (P < 0.001) such that is was decreased to ~ 1-4 pmol min^¬1^ in all species at 3°C. These differences were, however, not reflected in their respective Q_10_ values, where *D. montana* (3.5 ± 0.6) tended to have a higher Q_10_ than *D. melanogaster* (2.3 ± 0.2), but relatively similar to that of *D. birchii* (2.9 ± 0.6) (Fig. 2D).

Rates of Na^+^ reabsorption varied significantly between species at warm temperature (Fig. 2E) with values of 76.0 ± 6.9, 70.5 ± 6.2 and 56.0 ± 8.1 pmol min^−1^ for *D. montana, D. melanogaster* and *D. birchii*, respectively (P = 0.001). Cold exposure decreased Na^+^ reabsorption in all species (P < 0.001), and the absolute reduction was similar across the three species (no interaction, P = 0.552) such that the highest Na^+^ reabsorption was found in *D. montana* (25.4 ± 5.6 pmol min^−1^), followed by *D. melanogaster* (12.1 ± 2.6 pmol min^−1^), while *D. birchii* switched to net Na^+^ secretion (−7.4 ± 4.1 pmol min^−1^) (Fig. 2E). The negative rate of reabsorption found for *D. birchii* prevents a calculation of a Q_10_, however, when comparing Q_10_’s for the two other species we found a non-significant tendency (P = 0.096) for a lower Q_10_ in the cold tolerant *D. montana* (1.9 ± 0.2) compared to that of *D. melanogaster* (2.6 ± 0.2) (Fig. 2F).

The different proportional effects of temperature on Na^+^ and K^+^ reabsorption were also reflected in large and significant differences in the temperature effect ratio of Na^+^ vs. K^+^ reabsorption (F_2_,_18_ = 12.9, P < 0.001). Thus, *D. birchii* was characterized by a much larger reduction in Na^+^ reabsorption relative to the reduction in K^+^ reabsorption (temperature effect ratio of 11.7 ± 1.9) than *D. montana* (temperature effect ratio of 1.9 ± 0.5) with intermediate values for *D. melanogaster* (temperature effect ratio of 5.1 ± 1.4) (Fig. 4A). In other words, the cold sensitive species (*D. birchii*) reduces Na^+^ reabsorption rate ~ 12 fold more than it reduces K^+^ reabsorption rate while the tolerant species (*D. montana*) only reduces Na^+^ reabsorption rate 2 fold compared to the reduction in K^*+*^ reabsorption rate.

### Intraspecific differences in hindgut transport during cold exposure

In a parallel set of experiments we tested how acclimation of *D. melanogaster* to high (23°C), intermediate (19°C) and low (15°C) temperature affected hindgut reabsorption. Overall the effects of cold acclimation (Fig. 3) were very similar to those found for cold adaptation (Fig. 2). There was a significant difference in fluid reabsorption rate between acclimation groups (P = 0.050) which were largely driven by differences measured at warm temperature (Fig. 3A). Thus, fluid reabsorption rate in warm-acclimated *D. melanogaster* (1.2 ± 0.1 nL min^−1^) was higher than that of controls (19°C acclimated; 0.9 ± 0.1 nL min^−1^) and cold-acclimated flies (0.8 ± 0.1 nL min^−1^). Exposure to cold drastically lowered these rates (P < 0.001) in all acclimation groups but magnitude of this reduction was dependent on the acclimation treatment (P < 0.001). Cold-acclimated flies maintained a higher rate of fluid reabsorption at 3°C (0.24 ± 0.03 nL min^−1^) compared to control flies (0.15 ± 0.03 nL min^−1^), which in turn had higher reabsorption rates than their warm-acclimated conspecifics (0.11 ± 0.01 nL min^−1^). These differences also manifest in the derived Q_10_’s (P < 0.001) which ranged from 3.4 ± 0.3 in warm-acclimated *D*. *melanogaster* to 2.6 ± 0.3 in controls and 1.8 ± 0.1 in the cold-acclimated conspecifics (Fig. 3B).

**Figure 3.**
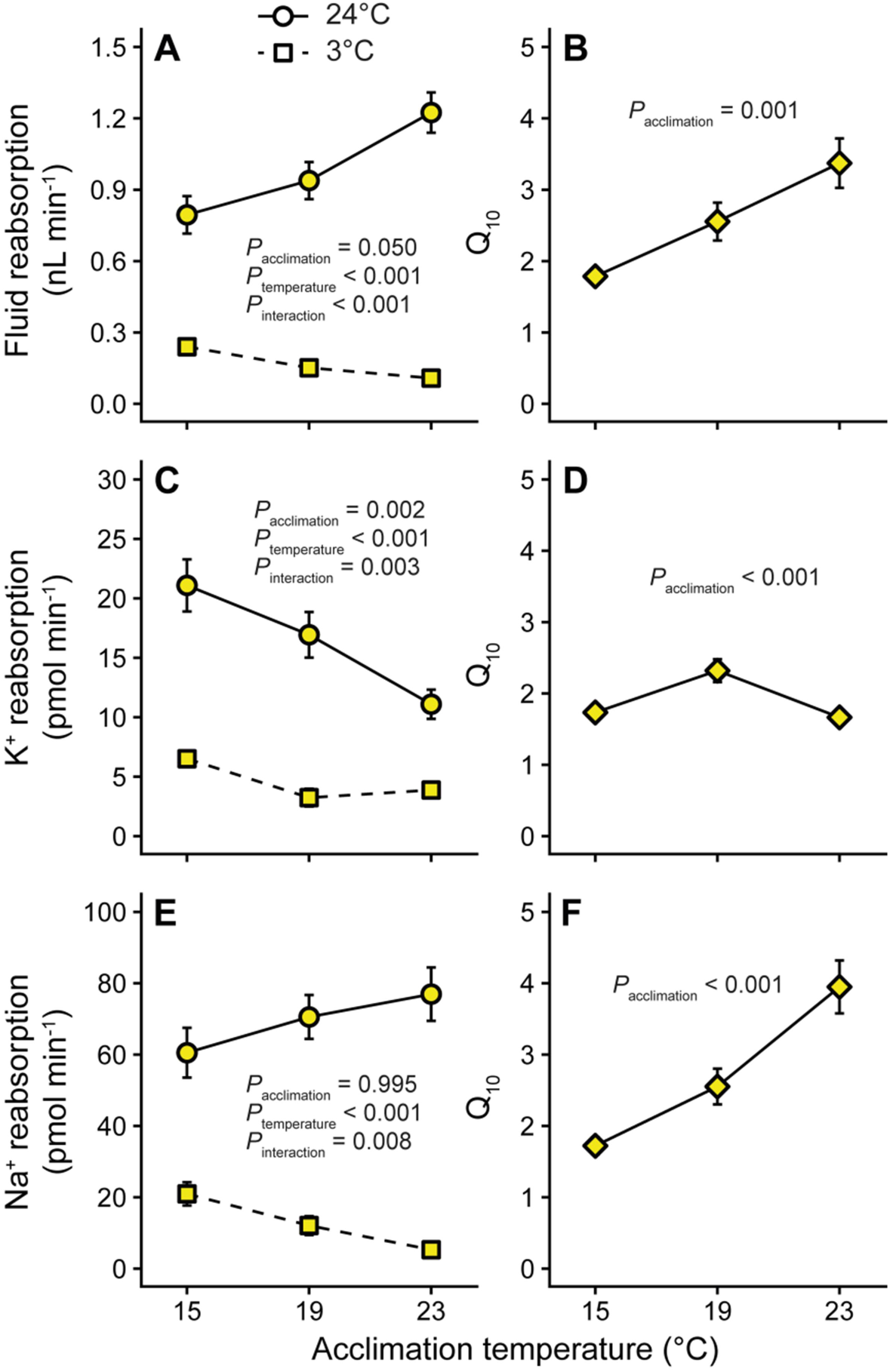
The effect of cold exposure on hindgut reabsorption in acclimated D. melanogaster. Reabsorption as the hindgut (left column) was measured at 24°C (circles) and 3°C (squares) in acclimated *D. melanogaster* and representative Q_10_’s were calculated (right column) for fluid (A and B), K^+^ (C and D), and Na^+^ (E and F) reabsorption. N = 9, 7, and 9 per acclimation temperature and temperature combination, and error bars not shown are covered by the symbols.

K^+^ reabsorption (Fig. 3C) was significantly influenced by acclimation temperature, P = 0.002) with the highest rates measured in the cold-acclimated flies (21.1 ± 2.2 pmol min^−1^), intermediate rates measured in control flies (16.9 ± 1.9 pmol min^−1^) and the lowest rates measured in warm-acclimated flies (11.1 ± 1.2 pmol min^−1^). These rates were markedly reduced by exposure to low temperature (P < 0.001), but the degree of suppression depended on acclimation temperature (F_2,22_ = 7.9, P = 0.003) such that cold-acclimated flies were better able to maintain rates of K^+^ reabsorption (6.5 ± 0.5 pmol min^−1^) compared to both control and warm-acclimated flies (3.2 ± 0.7 pmol min^−1^ and 3.9 ± 0.4 pmol min^−1^, respectively). These patterns resulted in Q_10_ being highest in control flies (2.3 ± 0.2), while Q_10_ was similar in the cold- and warm-acclimated conspecifics (1.7 ± 0.0 and 1.7 ± 0.1, respectively) (Fig. 3D).

Rates of Na^+^ reabsorption were generally independent of acclimation temperature (P = 0.995) but there was a highly significant interaction between acclimation group and the response to cooling (Fig. 3E). Thus, Na^+^ reabsorption was markedly reduced by exposure to 3°C (P < 0.001) but this reduction was dependent on acclimation temperature (P = 0.008) such that cold-acclimated flies had the highest rate of Na^+^ reabsorption at 3°C (21.0 ± 3.3 pmol min^−1^) while control flies reabsorbed 12.1 ± 2.6 pmol min^−1^ and warm-acclimated flies only managed to reabsorb 5.3 ± 1.1 pmol min^−1^. The derived Q_10_ therefore also showed significant differences between acclimation treatments (P < 0.001) with the lowest Q_10_ found in cold tolerant, cold-acclimated flies (1.7 ± 0.1), the highest in warm-acclimated flies (3.9 ± 0.4), and an intermediate Q_10_ in control flies (2.6 ± 0.2) (Fig. 3F).

To evaluate the cold induced reduction in Na^+^ reabsorption relative to the cold induced reduction in K^+^ reabsorption we calculated the temperature effect ratios of ΔNa^+^ reabsorption/ΔK^+^ reabsorption for the three acclimation groups. Here we found a clear effect of acclimation (Fig. 4B, P = 0.004) such that the cold-acclimated flies had the lowest ratio (2.8 ± 0.4), control flies an intermediate ratio (5.1 ± 1.4), and warm-acclimated flies the highest ratio (12.4 ± 2.8). Accordingly, warm acclimated flies have a much larger reduction in Na^+^ reabsorption rate than cold acclimated flies when this is calculated proportional to their reduction in K^+^ reabsorption rate.

**Figure 4.**
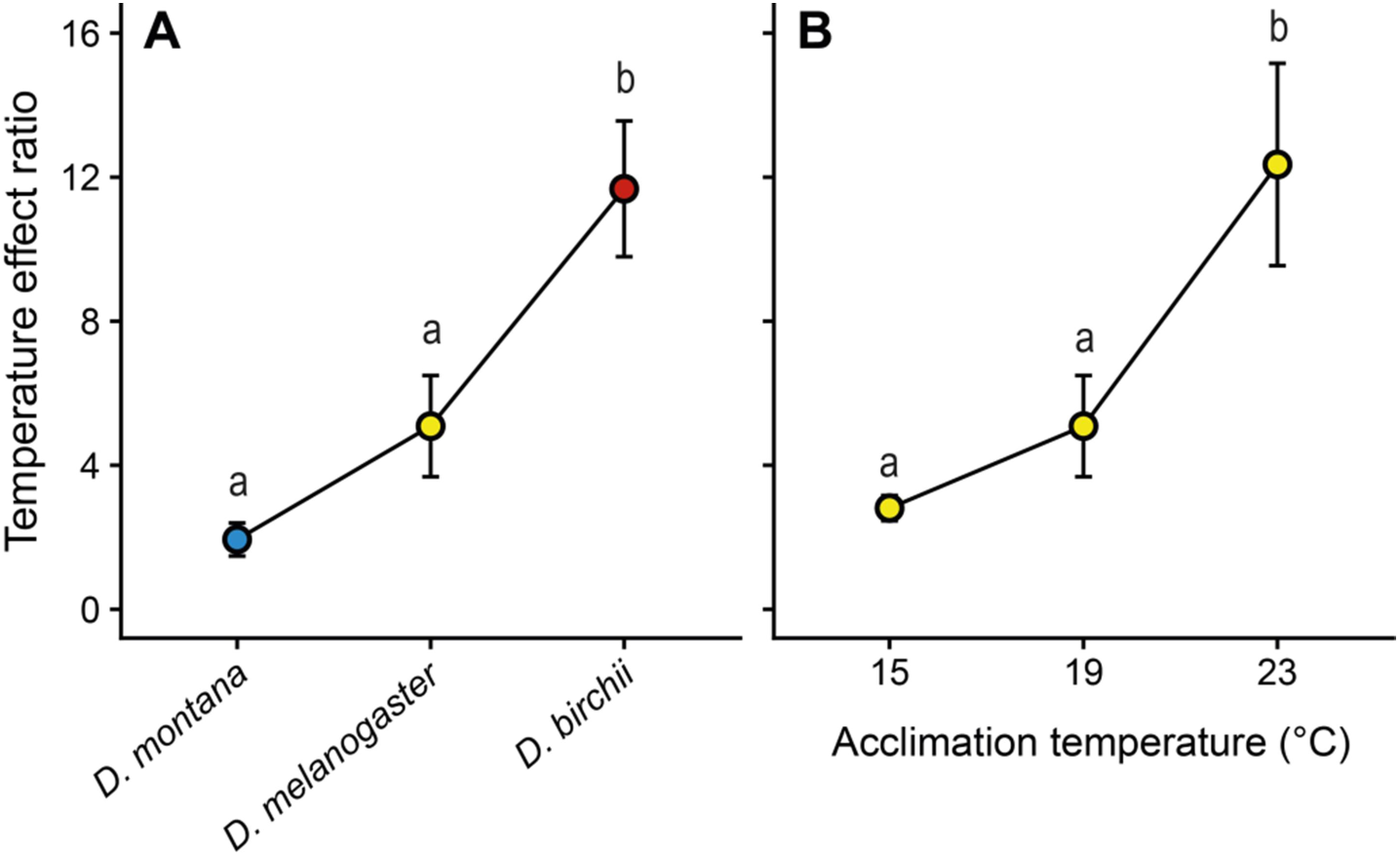
The temperature effect ratio between K^+^ and Na^+^ reabsorption. The temperature effect ratio is presented here as the absolute effect of cold exposure on Na^+^ reabsorption relative to the effect on K^+^ reabsorption in A) three *Drosophila* species and B) acclimated *D. melanogaster.* N = 7 per species and 9, 7, and 9 per acclimation temperature for *D*. *melanogaster* (note that the data for *D. melanogaster* has been used twice as these were acclimated to 19°C for the interspecific experiments). Groups not sharing letters are statistically different based on Tukey’s Honest Significant Differences *post hoc* tests.

## Discussion

### A novel assay to study epithelial function in the hindgut of Drosophila

Maintenance of extracellular volume and ion composition is essential to survival in all animals (Bernard, 1872; Beyenbach, 2016; Edney, 1977; Harrison et al., 2012) and in insects it is primarily the Malpighian tubules and the gut that are responsible for maintaining osmotic and ionic balance of the hemolymph (Edney, 1977; Phillips, 1970). Studies of bulk ion and water transport in small insects (<10 mg) has hitherto focused on Malpighian tubules because of the relative ease in preparing and measuring net secretion in this system (using the Ramsay assay, see Dow et al. (1994) and Rheault and O’Donnell (2004)). Here, we present a novel experimental assay which allows *in vitro* measurements of bulk fluid and ion movement across the hindgut epithelia in *Drosophila.* Fluid reabsorption of the hindgut remained stable over time (at least for three hours, Fig. 2) which allows for repeated measurements. Furthermore, we found the predicted effects of metabolic blockade and pharmacological intervention (Table 2) and exposure to low temperature (general reduction in transport with a Q_10_ ~2 to 5, Fig. 2 and 3) on the rates of reabsorption. Lastly, by comparing the absolute rates of reabsorption from this study with measurements of primary urine secretion from *D. melanogaster* (from MacMillan et al. (2015a)) we find that the Malpighian tubules secrete ~ 3.4 times more fluid, ~ 28.5 times more K^+^ and ~ 6 *%* more Na^+^ than is being reabsorbed. Some of these differences could relate to the methodological differences between the Ramsay assay and the hindgut assay presented here: In the Ramsay assay it is often difficult to include the lower, reabsorptive segment of the tubule (see O’Donnell and Maddrell (1995)) and there is no fluid pressure to be overcome at the end of the secreting tubule, meaning that secretion is likely overestimated. In the hindgut assay, the Malpighian tubules may work against a fluid pressure inside the gut, and the primary urine also passes through the reabsorbtive tubule segment before passing into the hindgut, such that there is less substrate (e.g. gut content and primary urine) to reabsorb from. It is also possible that there is some reabsorption from the mid gut and lastly, we did not quantify the amount of fluid and ions leaving the system as fecal droplets which could also explain why rates of secretion at the Malpighian tubules and absorption at the hindgut are not aligned. Sporadic measurements of the ion concentration of the fecal droplets of control *D. melanogaster* did, however, reveal a high K^+^ concentration (95.6 ± 6.5 mmol L^−1^, N = 14) and a moderate Na^+^ concentration (48.2 ± 3.5 mmol L^−1^, N = 14) which could at least partially account for the different transport rates observed. The dynamic interplay between secretion and reabsorption is also briefly demonstrated from the studies in which we inhibit ATP-dependent transport at the Malpighian tubules with NaCN. This treatment reduced both fluid and cation reabsorption at the hindgut to an extent similar to that of NaCN poisoning of the hindgut itself (Table 2). This effect is probably not caused by CN^−^ ions being transported by the fluid flow as NaCN blockade at both ends of the system caused even higher reductions in fluid, but it demonstrates logically that hindgut reabsorption is dependent on the provision from the osmoregulatory organs upstream. This was also seen from the experiment in which we decreasing buffer Na^+^ concentration. This treatment leads to a reduction in the primary urine Na^+^ concentration (Beyenbach, 2019; MacMillan et al., 2015a) and accordingly we find here that Na^+^ reabsorption is also reduced (Table 2).

The mechanisms underlying ion and water reabsorption in the insect hindgut have been most studied in locusts (primarily by Phillips and co-workers, see Black et al. (1987), Hanrahan and Phillips (1983), Phillips (1981), Phillips et al. (1987), and Phillips et al. (1996)). Using this model system, they proposed the involvement of basolateral and intermembrane (i.e. within the scalariform complex) Na^+^/K^+^ ATPase activity driving particularly Na^+^ and fluid reabsorption, apical V-type H^+^ ATPase activity coupled to H^+^ and Cl^−^ recycling driving Cl^−^ and secondarily K^+^ reabsorption, and the involvement of an unidentified apical Cl^−^ ATPase reabsorbing Cl^−^ combined with basolateral Cl^−^ channels and K^+^ following secondarily through channels. It is currently thought that similar mechanisms drive reabsorption in the *Drosophila* hindgut (Larsen et al., 2014; Phillips et al., 1996), and seen from Table 2 the findings of using pharmacological blockade suggests that the model is highly similar as; 1) inhibition of Na^+^/K^+^ ATPase activity with ouabain greatly reduced Na^+^ and fluid reabsorption, 2) blockade of V-type H^+^ ATPase activity with bafilomycin A_1_ reduced fluid and Na^+^ reabsorption suggesting a role for H^+^ recycling and likely also the apical membrane potential in reabsorption of fluid and Na^+^, and 3) blockade of Cl^−^ channels and transporters with DIDS almost completely abolished transport, suggesting that all cation transport is either linked directly to Cl^−^ transport or is reliant on the movement of Cl^−^ to maintain charge neutrality. Furthermore, we demonstrate that at least some of these processes are modulated by cAMP which greatly increased reabsorption of both fluid and cations, particularly K^+^. Increased intracellular concentrations of cAMP are a typical response to several forms of neuroendocrine stimulation (Coast et al., 2002; Dow et al., 2018; Phillips and Audsley, 1995). These findings show that the assay responds in general accordance with the current model of insect hindgut reabsorption but we stress that the mechanisms involved in this transport are complex and likely involve a range of channels, transporters and neuroendocrine factors (with more being identified, see for instance Luan et al. (2015)), which should be studied in further detail.

The assay presented here requires relatively modest experimental equipment (a microscope, a camera, a micromanipulator, and a steady hand) and the simple experimental approach of this assay will allow researchers to address many questions regarding epithelial function of a model insect (*Drosophila*) and probably also other small insects such as mosquitos. The assay is more time consuming, but has several advantages compared to the SIET assay used currently to study ion flux. These advantages are primarily that it is possible to measure bulk transport and that it is possible to measure all ions while measurements of Na^+^ is difficult using the SIET system and often requires a non-physiologically low saline Na^+^ concentration (Naikkhwah and O’Donnell, 2012). Thus, just like the development and optimization of the Ramsay assay paved the way for a multitude of mechanistic and comparative studies (Beyenbach et al., 2010; Dow and Davies, 2003; Dow et al., 2018; Halberg et al., 2015; Wheeler and Coast, 1990), this hindgut assay holds a potential for future research on epithelial function that could be aimed at unraveling 1) details of the physiological mechanism driving hindgut reabsorption, 2) the humoral/neuroendocrine mechanisms that regulate hindgut reabsorption, 3) the dynamic interplay between secretion at the Malpighian tubules and reabsorption at the hindgut, and 4) the intricacies of how osmoregulatury function is recruited in *Drosophila* during stressful challenges such are dehydration, extreme temperatures or diet and salt stress.

### Cold adapted species preserve osmoregulatory capacity at low temperatures

Exposure to stressful cold causes chill sensitive insect species to lose ion balance characterized by hemolymph hyperkalemia. This depolarizes excitable tissues and has been shown to directly cause damage and thereby be a probable cause of injury and mortality at low temperature (Bayley et al., 2018; Overgaard and MacMillan, 2017). According to this paradigm a number of studies have indicated a role of sustained osmoregulatory function at low temperature (Andersen et al., 2017c; Des Marteaux et al., 2017; Gerber and Overgaard, 2018; MacMillan et al., 2015a; Yi and Lee, 2005). These studies have implicated adaptations in regards to both secretory and absorptive function, but in the case of small insects such as *Drosophila* it has been complicated to study osmoregulatory adaptations in bulk transport at the hindgut.

In the present study we examined three *Drosophila* species that are known to differ markedly in their cold tolerance. We have previously used this comparative model system and the difference in cold tolerance is exemplified by the cold tolerant *D*. *montana* having a markedly lower LT_50_, a much quicker recovery following a standard cold exposure, and a much lower temperature threshold for the transition into chill coma (see Andersen et al. (2015) and Andersen and Overgaard (2019)). Consistent with our hypothesis, we found dramatic interspecific differences in hindgut responses to low temperature. All species reduced ion and fluid reabsorption when exposed to cold but importantly the reductions in reabsorption of fluid and specific ions differed among species such that the most cold tolerant species (*D. montana*) was able to defend Na^+^ and fluid reabsorption compared to the reduction in K^+^ reabsorption while the more cold sensitive allospecifics (e.g. *D. birchii*) reduced Na^+^ and fluid reabsorption more compared to K^+^ reabsorption rates (see Fig. 2 and 4A). These responses can all be viewed as being adaptive because they will to prevent the characteristic hyperkalemic condition that is linked to chill injury. Specifically, preservation of Na^+^ reabsorption helps prevent loss of Na^+^ balance which, along with the maintained rate of fluid reabsorption, will secure hemolymph volume. Hyperkalemia in insects is often attributed to the reduction of hemolymph volume that concentrates K^+^ remaining in the extracellular fluid (MacMillan and Sinclair, 2011; Olsson et al., 2016). Preservation of fluid reabsorption of the cold adapted species exposed to cold is therefore provisionary for normokalemia, and this is further aided by a marked reduction in K^+^ reabsorption. These results are corroborated by previous findings in the same species, namely that the same trends have been found by Andersen et al. (2017c) who used the SIET method to demonstrate that cold tolerant *Drosophila* species (e.g. *D. montana*) lower K^+^ reabsorption more than their cold sensitive allospecifics. The interspecific differences we observe in the hindgut response to cold are more or less opposite to the effects observed for changes in secretory function in the Malpighian tubules earlier (MacMillan et al., 2015a), and the responses therefore reinforce each other. Specifically for the cold tolerant *D. montana* it is observed that K^+^ secretion is relatively higher in the Malpighian tubules and relatively lower in the hindgut in response to cold – both responses will act to prevent hemolymph hyperkalemia. Similarly, fluid and particularly Na^+^ secretion are reduced in cold-exposed Malpighian tubules of *D*. *montana* and both fluid and Na^+^ reabsorption are defended better – again this will help preserve hemolymph volume and Na^+^ concentration, ultimately preventing hemolymph hyperkalemia.

In *D*. *birchii* we observed a reversal of Na^+^ flux from Na^+^ reabsorption to functional Na^+^ secretion in (Fig. 2E). Leak of Na^+^ down electrochemical gradients from the hemolymph into the gut was first proposed as a cause of reduced hemolymph volume by MacMillan and Sinclair (2011) who studied changes in volume and composition of hemolymph during cold exposure in the fall field cricket (*Gryllus pennsylvanicus*). Here they found that Na^+^ and fluid leaked towards the gut, and since then several other reports have implicated similar disturbances in other insects (Des Marteaux and Sinclair, 2016; Gerber and Overgaard, 2018; Koštál et al., 2007; Koštál et al., 2004; Koštál et al., 2006) including *Drosophila* (MacMillan et al., 2015a; MacMillan et al., 2015b; MacMillan et al., 2016). Additionally, leak assays have been used to investigate the passive movement of Na^+^ over the gut in insects using fluorescent markers and these studies have generally found that cold sensitive *Drosophila* are more susceptible to transepithelial leak at low temperature (Andersen et al., 2017c; MacMillan et al., 2017). Despite these observations, this is the first study to present direct measurements of cold-induced leak of ions down their electrochemical gradients across a transporting epithelium in a chill susceptible insect.

### Thermal acclimation mitigates loss of osmoregulatory function in the cold

Differences in cold tolerance are also found within species (e.g. intraspecific variation) and insects are highly plastic in their lower thermal limits if given time to acclimate or acclimatize to cold (Lee, 2012; Mellanby, 1954; Overgaard et al., 2011; Sinclair et al., 2003), *D. melanogaster* included (Colinet and Hoffmann, 2012; MacMillan et al., 2017). It is therefore not surprising that cold-acclimated or winter-acclimatized *Drosophila* are more cold tolerant and are able to defend their hemolymph volume and ion balance better than their warm-acclimated or summer-acclimatized conspecifics (MacMillan et al., 2015b; MacMillan et al., 2016). Accordingly, recent studies have found that cold-acclimated insects are better able to maintain osmoregulatory function when exposed to stressful cold (Gerber and Overgaard, 2018; Yerushalmi et al., 2018; Yi and Lee, 2005). These studies have also all demonstrated plasticity in the function of Malpighian tubules and the gut, but prior to this study no measurements of bulk fluid and ion movements have been made for the hindgut of small insects like *Drosophila*.

Here we used *D*. *melanogaster* acclimated to three temperatures known to elicit changes in cold tolerance (Bubliy et al., 2002; MacMillan et al., 2015d; Schou et al., 2017). Similarly to the experiments on interspecific variation (see Fig. 2) we found differences between groups in their hindgut reabsorption capacity at low temperature (Fig. 3). Overall exposure to low temperature reduced reabsorption regardless of acclimation temperature, but like with the cold-adapted *D*. *montana*, cold-acclimated *D*. *melanogaster* were generally better at maintaining transport rates when it comes to fluid reabsorption (Fig. 2A and 2B), a larger reduction in K^+^ reabsorption (Fig. 2C and 2D), and an improved maintenance of Na^+^ reabsorption (Fig. 2E and 2F). These findings corroborate previous findings made in an orthopteran insect (Gerber and Overgaard, 2018), and further so by SIET measurements made on the hindgut of acclimated *D. melanogaster* (Yerushalmi et al., 2018). Combined with measurements of ion and fluid secretion by the Malpighian tubules of acclimated *D. melanogaster* (see Yerushalmi et al. (2018)), the overall result is highly similar to that described between species: improved cold tolerance is achieved through modulations to the osmoregulatory systems that act to protect hemolymph volume and Na^+^ concentration and alleviate hemolymph normokalemia.

## Conclusion

Overall there is now surmounting evidence that improved chill tolerance in *Drosophila* is tightly linked to the ability to better maintain osmoregulatory function at low temperature in both sides of the osmoregulatory system: The combined actions of the Malpighian tubules and the hindgut act in synchrony to prevent loss of hemolymph volume and hyperkalemia. This is achieved by 1) a greater reduction in K^+^ reabsorption and a better maintained K^+^ secretion, and 2) the maintenance of a higher rate of Na^+^ reabsorption and a lower relative rate of Na^+^ secretion which act to prevent loss of hemolymph volume by counteracting the passive leak of Na^+^ and water into the gut. However, while the present study identifies clear inter- and intraspecific differences in ion and fluid transport and their response to cold, it does not reveal the physiological mechanism behind these differences. Recent studies have suggested that neuroendocrine regulation plays a central role in modulating cold tolerance *via* actions on secretory mechanisms (see Terhzaz et al. (2015) and MacMillan et al. (2018)). However, in this and previous studies have demonstrated differences in osmoregulatory capacity are found in *in vitro* assays that are devoid of neuroendocrine input. It is therefore clear that some of the differences found in relation to acclimation and adaptation must be related to more permanent physiological adjustments. Such adjustments could include changes in protein expression or changes in membrane phospholipid composition which could affect the active transport capacity as well as the passive conductance of the epithelia.

## Data accessibility

All data is attached as supplementary files for review purposes, and the same files will be uploaded to the Dryad data repository should the article be accepted and published.

## Acknowledgements

The authors would like to thank Kirsten Kromand for help with fly and lab maintenance and Thomas Holm Pedersen for use of electrode puller.

## Author contributions

The study was conceived and designed by both authors. MKA performed the experiments, analyzed the data, and wrote first draft. Both authors contributed to and approved of the final version of the manuscript.

## References

Andersen, J. L., Manenti, T., Sørensen, J. G., MacMillan, H. A., Loeschcke, V. and Overgaard, J. (2015). How to assess Drosophila cold tolerance: chill coma temperature and lower lethal temperature are the best predictors of cold distribution limits. Functional Ecology 29, 55–65.

Andersen, M. K., Folkersen, R., MacMillan, H. A. and Overgaard, J. (2017a). Cold acclimation improves chill tolerance in the migratory locust through preservation of ion balance and membrane potential. Journal of Experimental Biology 220, 487–496.

Andersen, M. K., Jensen, S. O. and Overgaard, J. (2017b). Physiological correlates of chill susceptibility in Lepidoptera. Journal of insect physiology 98, 317–326.

Andersen, M. K., MacMillan, H. A., Donini, A. and Overgaard, J. (2017c). Cold tolerance of Drosophila species is tightly linked to epithelial K^+^ transport capacity of the Malpighian tubules and rectal pads. Journal of Experimental Biology, jeb. 168518.

Andersen, M. K. and Overgaard, J. (2019). The central nervous system and muscular system play different roles for chill coma onset and recovery in insects. Comparative Biochemistry and Physiology Part A: Molecular & Integrative Physiology 233, 10–16.

Bayley, J. S., Winther, C. B., Andersen, M. K., Grønkjær, C., Nielsen, O. B., Pedersen, T. H. and Overgaard, J. (2018). Cold exposure causes cell death by depolarization-mediated Ca^2+^ overload in a chill-susceptible insect. Proceedings of the National Academy of Sciences, 201813532.

Bernard, C. (1872). De la physiologie générale: Hachette.

Beyenbach, K. W. (2016). The plasticity of extracellular fluid homeostasis in insects. Journal of Experimental Biology 219, 2596–2607.

Beyenbach, K. W. (2019). Voltages and resistances of the anterior Malpighian tubule of Drosophila melanogaster. Journal of Experimental Biology 222, jeb201574.

Beyenbach, K. W. and Piermarini, P. M. (2008). Osmotic and ionic regulation in insects. Osmotic and ionic regulation: cells and animals, 231–293.

Beyenbach, K. W. and Piermarini, P. M. (2011). Transcellular and paracellular pathways of transepithelial fluid secretion in Malpighian (renal) tubules of the yellow fever mosquito Aedes aegypti. Acta Physiologica 202, 387–407.

Beyenbach, K. W., Skaer, H. and Dow, J. A. (2010). The developmental, molecular, and transport biology of Malpighian tubules. Annual review of entomology 55, 351–374.

Black, K., Meredith, J., Thomson, B., Phillips, J. and Dietz, T. (1987). Mechanisms and properties of sodium transport in locust rectum. Canadian Journal of Zoology 65, 3084–3092.

Bubliy, O. A., Riihimaa, A., Norry, F. M. and Loeschcke, V. (2002). Variation in resistance and acclimation to low-temperature stress among three geographical strains of Drosophila melanogaster. Journal of thermal biology 27, 337–344.

Coast, G. M., Orchard, I., Phillips, J. E. and Schooley, D. A. (2002). Insect diuretic and antidiuretic hormones.

Colinet, H. and Hoffmann, A. A. (2012). Comparing phenotypic effects and molecular correlates of developmental, gradual and rapid cold acclimation responses in Drosophila melanogaster. Functional Ecology 26, 84–93.

D’Silva, N. M., Donini, A. and O’Donnell, M. J. (2017). The roles of V-type H^+^-ATPase and Na^+^/K^+^-ATPase in energizing K^+^ and H^+^ transport in larval Drosophila gut epithelia. Journal of insect physiology 98, 284–290.

Davies, S.-A., Cabrero, P., Marley, R., Corrales, G. M., Ghimire, S., Dornan, A. J. and Dow, J. A. (2019). Epithelial Function in the Drosophila Malpighian Tubule: An In Vivo Renal Model. In Kidney Organogenesis, pp. 203–221: Springer.

Des Marteaux, L. E., Khazraeenia, S., Yerushalmi, G. Y., Donini, A., Li, N. G. and Sinclair, B. J. (2017). The effect of cold acclimation on active ion transport in cricket ionoregulatory tissues. Comparative Biochemistry and Physiology Part A: Molecular & Integrative Physiology.

Des Marteaux, L. E. and Sinclair, B. J. (2016). Ion and water balance in Gryllus crickets during the first twelve hours of cold exposure. Journal of insect physiology 89, 19–27.

Donini, A. and O’Donnell, M. J. (2005). Analysis of Na^+^, Cl^−^, K^+^, H^+^ and NH_4_^+^ concentration gradients adjacent to the surface of anal papillae of the mosquito Aedes aegypti: application of self-referencing ion-selective microelectrodes. Journal of Experimental Biology 208, 603–610.

Dow, J., Maddrell, S., Görtz, A., Skaer, N., Brogan, S. and Kaiser, K. (1994). The malpighian tubules of Drosophila melanogaster: a novel phenotype for studies of fluid secretion and its control. Journal of Experimental Biology 197, 421–428.

Dow, J. A. and Davies, S. A. (2003). Integrative physiology and functional genomics of epithelial function in a genetic model organism. Physiological reviews 83, 687–729.

Dow, J. A., Halberg, K. A., Terhzaz, S. and Davies, S. A. (2018). Drosophila as a Model for Neuroendocrine Control of Renal Homeostasis. Model Animals in Neuroendocrinology: From Worm to Mouse to Man, 81–100.

Edney, E. B. (1977). Excretion and Osmoregulation. In Water Balance in Land Arthropods, pp. 96–171: Springer.

Gerber, L. and Overgaard, J. (2018). Cold tolerance is linked to osmoregulatory function of the hindgut in Locusta migratoria. Journal of Experimental Biology, jeb. 173930.

Gerencser, G. A. and Zhang, J. (2003). Chloride ATPase pumps in nature: do they exist? Biological Reviews 78, 197–218.

Gupta, B. L., Wall, B. J., Oschman, J. L. and Hall, T. A. (1980). Direct microprobe evidence of local concentration gradients and recycling of electrolytes during fluid absorption in the rectal papillae of Calliphora. Journal of Experimental Biology 88, 21–48.

Halberg, K. A., Terhzaz, S., Cabrero, P., Davies, S. A. and Dow, J. A. (2015). Tracing the evolutionary origins of insect renal function. Nature Communications 6, 6800.

Hanrahan, J., Meredith, J., Phillips, J. and Brandys, D. (1984). Methods for the study of transport and control in insect hindgut. In Measurement of Ion Transport and Metabolic Rate in Insects, pp. 19–67: Springer.

Hanrahan, J. and Phillips, J. (1983). Cellular mechanisms and control of KCl absorption in insect hindgut. Journal of Experimental Biology 106, 71–89.

Harrison, J. F., Woods, H. A. and Roberts, S. P. (2012). Ecological and environmental physiology of insects: Oxford University Press.

Kellermann, V., Loeschcke, V., Hoffmann, A., Kristensen, T., Flojgaard, C., David, J., Svenning, J.-C. and Overgaard, J. (2012). Phylogenetic constraints in key functional traits behind species’ climate niches: patterns of desiccation and cold resistance across 95 Drosophila species. Evolution; international journal of organic evolution [P] 66, 3377–3389.

Koštál, V., Renault, D., Mehrabianova, A. and Bastl, J. (2007). Insect cold tolerance and repair of chill-injury at fluctuating thermal regimes: role of ion homeostasis. Comparative Biochemistry and Physiology Part A: Molecular & Integrative Physiology 147, 231–238.

Koštál, V., Vambera, J. and Bastl, J. (2004). On the nature of pre-freeze mortality in insects: water balance, ion homeostasis and energy charge in the adults of Pyrrhocoris apterus. Journal of Experimental Biology 207, 1509–1521.

Koštál, V., Yanagimoto, M. and Bastl, J. (2006). Chilling-injury and disturbance of ion homeostasis in the coxal muscle of the tropical cockroach (Nauphoeta cinerea). Comparative Biochemistry and Physiology Part B: Biochemistry and Molecular Biology 143, 171–179.

Larsen, E. H., Deaton, L. E., Onken, H., O’Donnell, M., Grosell, M., Dantzler, W. H. and Weihrauch, D. (2014). Osmoregulation and excretion. Comprehensive physiology 4, 405–573.

Lee, R. (2012). Insects at low temperature: Springer Science & Business Media.

Luan, Z., Quigley, C. and Li, H.-S. (2015). The putative Na^+^/Cl−-dependent neurotransmitter/osmolyte transporter inebriated in the Drosophila hindgut is essential for the maintenance of systemic water homeostasis. Scientific reports 5, 7993.

MacMillan, H. A., Andersen, J. L., Davies, S. A. and Overgaard, J. (2015a). The capacity to maintain ion and water homeostasis underlies interspecific variation in Drosophila cold tolerance. Scientific reports 5.

MacMillan, H. A., Andersen, J. L., Loeschcke, V. and Overgaard, J. (2015b). Sodium distribution predicts the chill tolerance of Drosophila melanogaster raised in different thermal conditions. American Journal of Physiology-Regulatory, Integrative and Comparative Physiology 308, R823–R831.

MacMillan, H. A., Baatrup, E. and Overgaard, J. (2015c). Concurrent effects of cold and hyperkalaemia cause insect chilling injury. In Proc. R. Soc. B, vol. 282, pp. 20151483: The Royal Society.

MacMillan, H. A., Ferguson, L. V., Nicolai, A., Donini, A., Staples, J. F. and Sinclair, B. J. (2015d). Parallel ionoregulatory adjustments underlie phenotypic plasticity and evolution of Drosophila cold tolerance. Journal of Experimental Biology 218, 423–432.

MacMillan, H. A., Nazal, B., Wali, S., Yerushalmi, G. Y., Misyura, L., Donini, A. and Paluzzi, J.-P. (2018). Anti-diuretic activity of a CAPA neuropeptide can compromise Drosophila chill tolerance. Journal of Experimental Biology, jeb. 185884.

MacMillan, H. A., Schou, M. F., Kristensen, T. N. and Overgaard, J. (2016). Preservation of potassium balance is strongly associated with insect cold tolerance in the field: a seasonal study of Drosophila subobscura. Biology letters 12, 20160123.

MacMillan, H. A. and Sinclair, B. J. (2011). The role of the gut in insect chilling injury: cold-induced disruption of osmoregulation in the fall field cricket, Gryllus pennsylvanicus. The Journal of Experimental Biology 214, 726–734.

MacMillan, H. A., Yerushalmi, G. Y., Jonusaite, S., Kelly, S. P. and Donini, A. (2017). Thermal acclimation mitigates cold-induced paracellular leak from the Drosophila gut. Scientific reports 7, 8807.

Mellanby, K. (1954). Acclimatization and the thermal death point in insects.

Naikkhwah, W. and O’Donnell, M. J. (2012). Phenotypic plasticity in response to dietary salt stress: Na^+^ and K^+^ transport by the gut of Drosophila melanogaster larvae. Journal of Experimental Biology 215, 461–470.

O’Donnell, M. J., Ianowski, J. P., Linton, S. M. and Rheault, M. R. (2003). Inorganic and organic anion transport by insect renal epithelia. Biochimica et Biophysica Acta (BBA)-Biomembranes 1618, 194–206.

O’Donnell, M. J. and Maddrell, S. (1995). Fluid reabsorption and ion transport by the lower Malpighian tubules of adult female Drosophila. Journal of Experimental Biology 198, 1647–1653.

Olsson, T., MacMillan, H. A., Nyberg, N., Stærk, D., Malmendal, A. and Overgaard, J. (2016). Hemolymph metabolites and osmolality are tightly linked to cold tolerance of Drosophila species: a comparative study. Journal of Experimental Biology 219, 2504–2513.

Overgaard, J., Kristensen, T. N., Mitchell, K. A. and Hoffmann, A. A. (2011). Thermal tolerance in widespread and tropical Drosophila species: does phenotypic plasticity increase with latitude? The American Naturalist 178, S80–S96.

Overgaard, J. and MacMillan, H. A. (2017). The Integrative Physiology of Insect Chill Tolerance. Annual Review of Physiology 79, null.

Phillips, J. (1981). Comparative physiology of insect renal function. American Journal of Physiology-Regulatory, Integrative and Comparative Physiology 241, R241–R257.

Phillips, J. and Audsley, N. (1995). Neuropeptide control of ion and fluid transport across locust hindgut. American Zoologist 35, 503–514.

Phillips, J., Hanrahan, J., Chamberlin, M. and Thomson, B. (1987). Mechanisms and control of reabsorption in insect hindgut. Advances in insect physiology 19, 329–422.

Phillips, J., Wiens, C., Audsley, N., Jeffs, L., Bilgen, T. and Meredith, J. (1996). Nature and control of chloride transport in insect absorptive epithelia. Journal of Experimental Zoology Part A: Ecological Genetics and Physiology 275, 292–299.

Phillips, J. E. (1970). Apparent transport of water by insect excretory systems. American Zoologist 10, 413–436.

R Core Team. (2019). R: A language and environment for statistical computing. R Foundation for Statistical Computing: See http://www.R-project.org/.

Ramsay, J. (1954). Active transport of water by the Malpighian tubules of the stick insect, Dixippus morosus (Orthoptera, Phasmidae). Journal of Experimental Biology 31, 104–113.

Rheault, M. R. and O’Donnell, M. J. (2004). Organic cation transport by Malpighian tubules of Drosophila melanogaster: application of two novel electrophysiological methods. Journal of Experimental Biology 207, 2173–2184.

Schindelin, J., Rueden, C. T., Hiner, M. C. and Eliceiri, K. W. (2015). The ImageJ ecosystem: An open platform for biomedical image analysis. Molecular reproduction and development 82, 518–529.

Schou, M. F., Mouridsen, M. B., Sørensen, J. G. and Loeschcke, V. (2017). Linear reaction norms of thermal limits in Drosophila: predictable plasticity in cold but not in heat tolerance. Functional Ecology 31, 934–945.

Sinclair, B. J., Vernon, P., Klok, C. J. and Chown, S. L. (2003). Insects at low temperatures: an ecological perspective. Trends in ecology & evolution 18, 257–262.

Terhzaz, S., Teets, N. M., Cabrero, P., Henderson, L., Ritchie, M. G., Nachman, R. J., Dow, J. A., Denlinger, D. L. and Davies, S.-A. (2015). Insect capa neuropeptides impact desiccation and cold tolerance. Proceedings of the National Academy of Sciences 112, 2882–2887.

Wall, B. and Oschman, J. (1975). Structure and function of the rectum in insects. Fortschr. Zool 23, 193–222.

Wall, B. J. and Oschman, J. L. (1970). Water and solute uptake by rectal pads of Periplaneta americana. American Journal of Physiology 218, 1208–1215.

Wheeler, C. H. and Coast, G. M. (1990). Assay and characterisation of diuretic factors in insects. Journal of insect physiology 36, 23–34.

Yerushalmi, G. Y., Misyura, L., MacMillan, H. A. and Donini, A. (2018). Functional plasticity of the gut and the Malpighian tubules underlies cold acclimation and mitigates cold-induced hyperkalemia in Drosophila melanogaster. The Journal of Experimental Biology 221.

Yi, S.-X. and Lee, R. E. (2005). Changes in gut and Malpighian tubule transport during seasonal acclimatization and freezing in the gall fly Eurosta solidaginis. Journal of Experimental Biology 208, 1895–1904.

